# Substrate Affinity and Specificity of the *Sc*Sth1p Bromodomain Are Fine-Tuned for Versatile Histone Recognition

**DOI:** 10.1101/559005

**Authors:** Bartlomiej J. Blus, Hideharu Hashimoto, Hyuk-Soo Seo, Aleksandra Krolak, Erik W. Debler

**Author notes:** These authors contributed equally (Bartlomiej J. Blus, Hideharu Hashimoto). Correspondence (B.J.B.) and (E.W.D).

## Abstract

Bromodomains recognize a wide range of acetylated lysine residues in histones and other nuclear proteins. Substrate specificity is critical for their biological function and arises from unique acetyl-lysine binding sites formed by variable loop regions. Here, we analyzed substrate affinity and specificity of the yeast *Sc*Sth1p bromodomain, an essential component of the “<u>R</u>emodels the <u>S</u>tructure of <u>C</u>hromatin” complex, and found that the wild-type bromodomain preferentially recognizes H3K14ac and H4K20ac peptides. Mutagenesis studies—guided by our crystal structure determined at 2.7 Å resolution—revealed loop residues Ser1276 and Trp1338 as key determinants for such interactions. Strikingly, point mutations of each of these residues substantially increased peptide binding affinity and selectivity, respectively. Our data demonstrate that the *Sc*Sth1p bromodomain is not optimized for binding to an individual acetylation mark, but fine-tuned for interactions with several such modifications, consistent with the versatile and multivalent nature of histone recognition by reader modules such as bromodomains.

**Highlights:**

- The *Sc*Sth1p bromodomain preferentially recognizes H3K14ac and H4K20ac peptides
- Ser1276 and Trp1338 are key determinants of substrate affinity and specificity
- Mutations of these residues drastically increase substrate affinity and specificity
- The *Sc*Sth1p bromodomain is fine-tuned for promiscuous histone tail recognition

Graphical Abstract

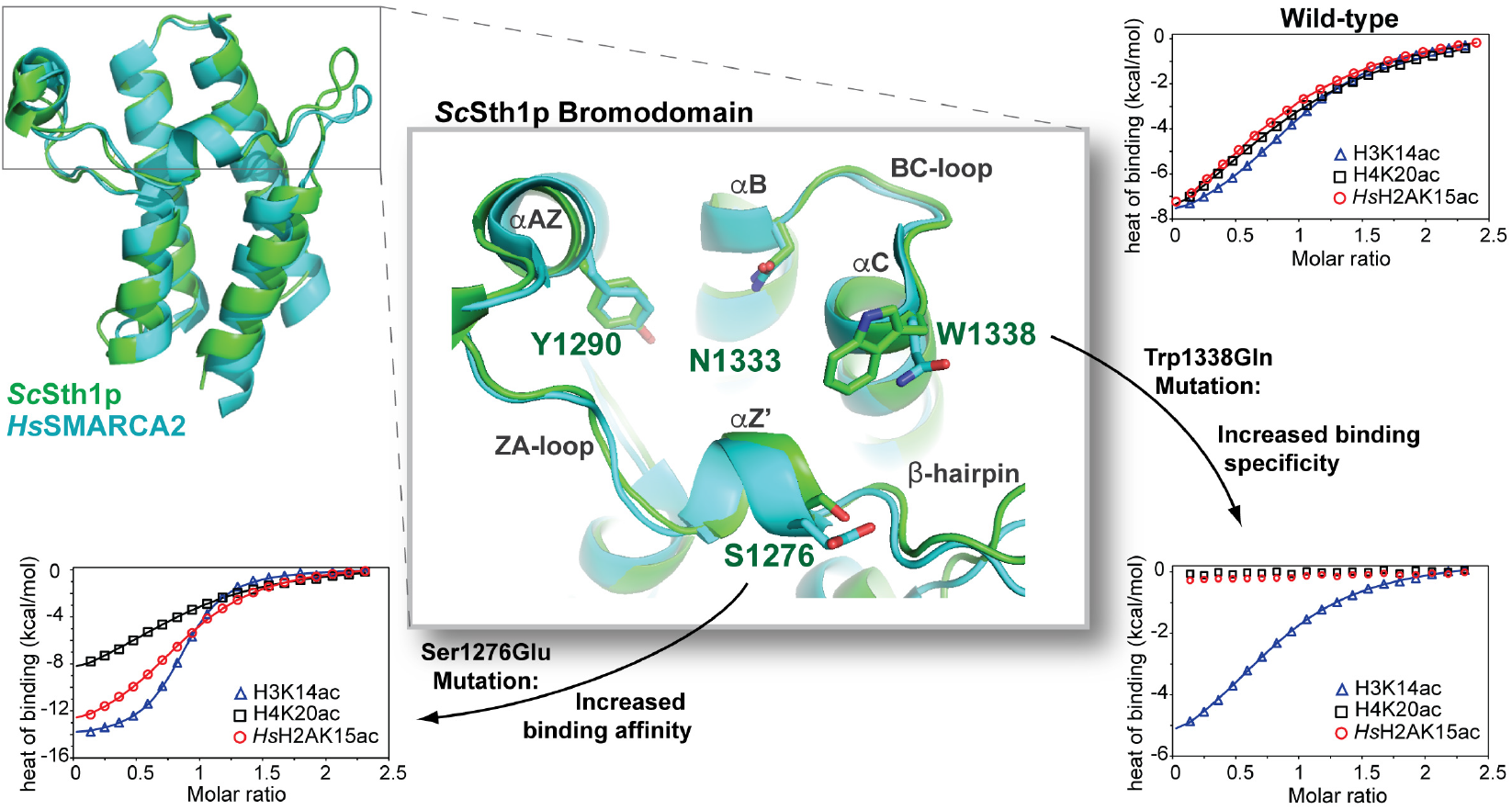

## Introduction

Protein acetylation is a ubiquitous post-translational modification primarily found in eukaryotes. Site-specific acetylation of lysine residues in histone tails plays a major role in regulating chromatin structure and gene expression (<u>Garcia et al., 2007</u>; <u>Huang et al., 2015</u>; <u>Latham and Dent, 2007</u>). Acetylated histone tails are a hallmark of active genes and multiple acetyl-lysine marks are often enriched at active promoter transcription start sites (TSSs) (<u>Garcia et al., 2007</u>). Histone acetylation is a dynamic and reversible process mediated by histone acetyltransferases and deacetylases (<u>Weiner et al., 2015</u>). The acetyl-lysine is recognized primarily by bromodomains, but also by the YEATS and Tudor domains (<u>Filippakopoulos et al., 2012</u>; <u>Jurkowska et al., 2017</u>; <u>Zhao et al., 2017</u>). Besides acetylation, bromodomains have been shown to recognize propionylated and butyrylated lysines (<u>Vollmuth and Geyer, 2010</u>). Consistent with their central role in chromatin biology and gene transcription, many bromodomains become dysregulated in numerous diseases including cancer and inflammatory disorders, and thus have emerged as prominent epigenetic drug targets (<u>Filippakopoulos and Knapp, 2014</u>; <u>Fujisawa and Filippakopoulos, 2017</u>).

Bromodomains consist of a left-handed four-helical bundle (αZ, αA, αB, αC) and variable ZA and BC loops involved in substrate recognition (<u>Dhalluin et al., 1999</u>; <u>Filippakopoulos et al., 2012</u>). They have conserved motifs in their helices and loops and can be divided into eight families using a structure-based alignment (<u>Filippakopoulos and Knapp, 2012</u>; <u>Filippakopoulos et al., 2012</u>). Most bromodomains contain PDY (Pro-Asp-Tyr) and P*ϕ*D (*ϕ* denotes a hydrophobic residue) motifs in the ZA-loop. Other bromodomain features are conserved to a lesser extent and are family-specific. For instance, families I and II contain the WPF (Trp-Pro-Phe) “shelf” motif within the ZA loop, which is equivalent to the E/D*ϕ*F motif in family VIII. The acetyl-lysine residue is recognized by a conserved hydrophobic pocket located at the end of the four-helix bundle in each bromodomain (<u>Dhalluin et al., 1999</u>). In particular, two highly conserved residues—an Asn at the C-terminus of the αB helix and a Tyr in the ZA loop—form canonical hydrogen bonds with the acetyl moiety in the target protein or peptide and the conserved network of water molecules in the acetyl-lysine binding pocket, respectively. In stark contrast to this conserved pocket, the surrounding surface area is formed by highly variable regions which confer sequence-specific recognition of the residues adjacent to the acetylated lysine (<u>Vidler et al., 2012</u>). Biochemical profiling of histone peptide interactions has revealed that most bromodomains recognize several different acetyl-lysine marks with micromolar to millimolar affinities (<u>Filippakopoulos and Knapp, 2012</u>). In addition to this binding promiscuity, multivalency of histone recognition by more than one “reader” module is well established, generating avidity and rationalizing why high affinity of recognition motifs to individual marks is often not observed (<u>Ruthenburg et al., 2007</u>).

In order to examine the interplay between substrate binding affinity and specificity, we set out to characterize these properties in the *Sc*Sth1p bromodomain from *Saccharomyces cerevisiae*. This bromodomain is among the few yeast bromodomains that bind distinct acetylated histone peptides with apparent high affinity (<u>Zhang et al., 2010</u>). The *Sc*Sth1p protein is an ATP-dependent helicase associated with the “<u>R</u>emodels the <u>S</u>tructure of <u>C</u>hromatin” (RSC) complex involved in transcription regulation and nucleosome positioning (<u>Cairns et al., 1996</u>; <u>Lorch et al., 2018</u>; <u>Rawal et al., 2018</u>). The C-terminal bromodomain in *Sc*Sth1p is essential and recruits the RSC complex to active TSSs (<u>Du et al., 1998</u>). In this study, we determined the affinity of the *Sc*Sth1p bromodomain towards several acetylated histone peptides by isothermal titration calorimetry (ITC) and report its crystal structure at 2.7 Å resolution. Comparison with other bromodomains allowed us to identify loop residues that are unique to *Sc*Sth1p. Strikingly, point mutations of these residues substantially increase substrate specificity and binding affinity of the *Sc*Sth1p bromodomain, corroborating the notion that bromodomains are fine-tuned for versatile histone recognition.

## Results

### The *Sc*Sth1p Bromodomain Preferentially Recognizes H3K14ac and H4K20ac Peptides

Using the SMART database (<u>Letunic et al., 2015</u>), we designed a *Sc*Sth1p bromodomain construct encoding residues 1250-1359, which was expressed as a stable protein for biophysical and structural characterization (Figures 1A and 1B). The *Sc*Sth1p bromodomain was previously shown to preferentially bind to three acetylated peptides derived from human histones: H3K14ac, H4K20ac, and *Hs*H2AK15ac (<u>Zhang et al., 2010</u>). Using isothermal titration calorimetry (ITC), we found that the *Sc*Sth1p bromodomain binds these peptides with dissociation constants (K_d_s) of ∼18 µM, 48 µM, and 44 µM, respectively (Figure 2B). These interactions are acetyl-lysine specific, as the unmodified histone peptides did not bind (Figures S1A and S1B). Likewise, alanine mutations of the two highly conserved acetyl-lysine recognizing residues, Tyr1290Ala and Asn1333Ala, completely abolished the *Sc*Sth1p bromodomain interactions with these acetylated peptides (Figures S1C and S1D). While the human and yeast H3 and H4 histone tails are identical (which is why we omit their species designations in this manuscript), the histone H2A sequences vary significantly between the two species. Due to these differences, we also tested two yeast peptides homologous to *Hs*H2AK15ac: *Sc*H2AK13ac and *Sc*Htz1K14ac (Figure 2A). Interestingly, these two peptides do not interact with the *Sc*Sth1p bromodomain (Figure 2C), suggesting that histone residues adjacent to the acetyl-lysine are critical for binding.

**Figure 1.**
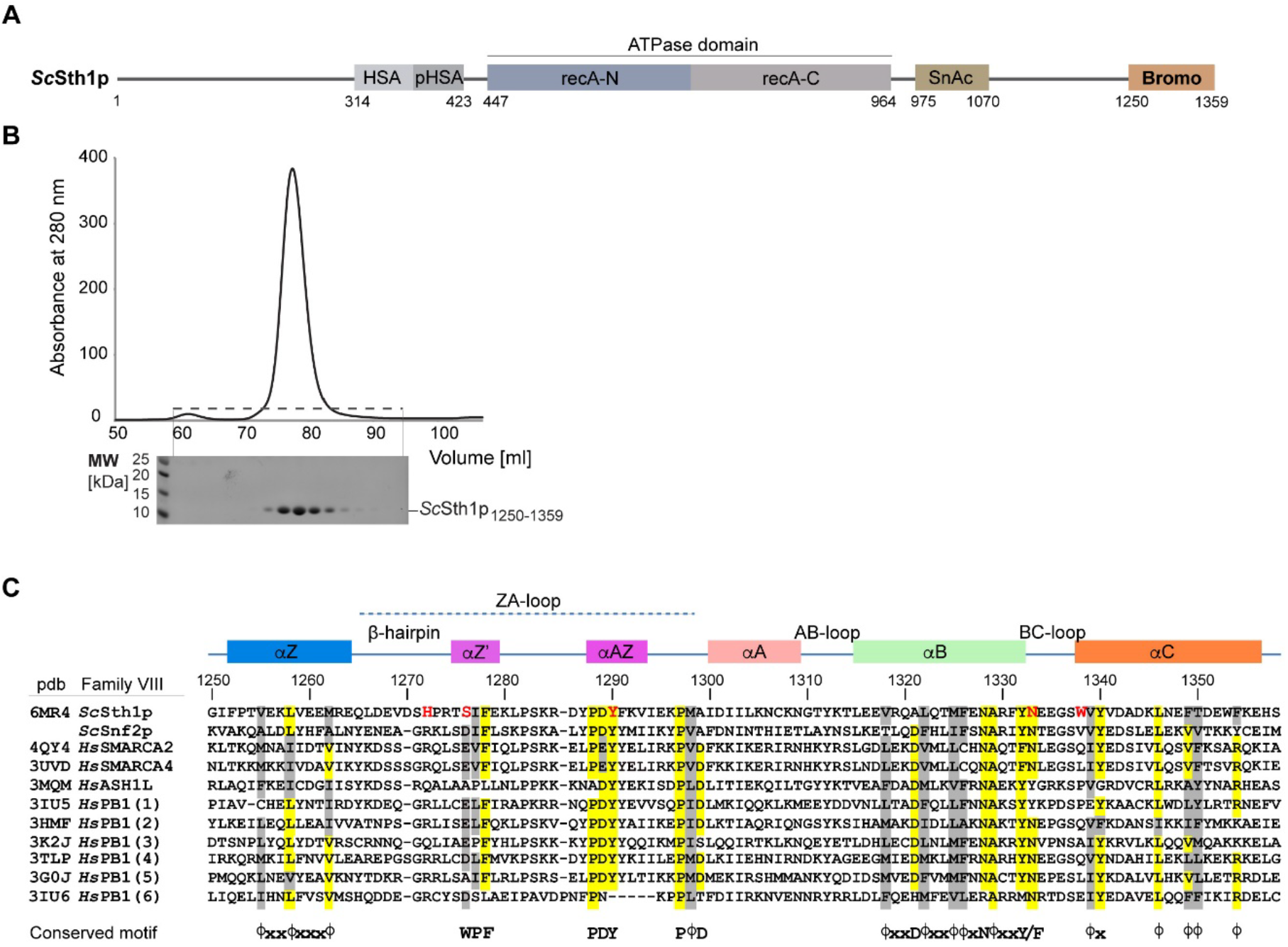
Purification and Sequence Alignment of the *Sc*Sth1p bromodomain. (A) Domain architecture of the *Sc*Sth1p protein. The N-terminal helicase/SANT-associated (HSA) domain, the catalytic ATPase module consisting of the recA domains, Snf2 ATP coupling (SnAc) domain, and the C-terminal bromodomain are shown. (B) A size-exclusion chromatography (SEC) elution profile for the *Sc*Sth1p bromodomain (residues 1250-1359) is shown together with peak elution fractions analyzed by SDS-PAGE. The bromodomain elutes as a single monomeric peak. (C) A structure-guided multiple sequence alignment of the *Sc*Sth1p bromodomain with similar yeast and human bromodomains. The sequences of the six bromodomains in the human PB1 protein are displayed. Secondary structure elements and the loop regions are shown above the alignment. Similar and identical residues within the conserved bromodomain sequences are colored grey and yellow, respectively, and the residues mutated in this study are highlighted in red. Highly-conserved sequence motifs across all bromodomain families are shown below the alignment. Φ and x denote hydrophobic and any residues, respectively.

**Figure 2.**
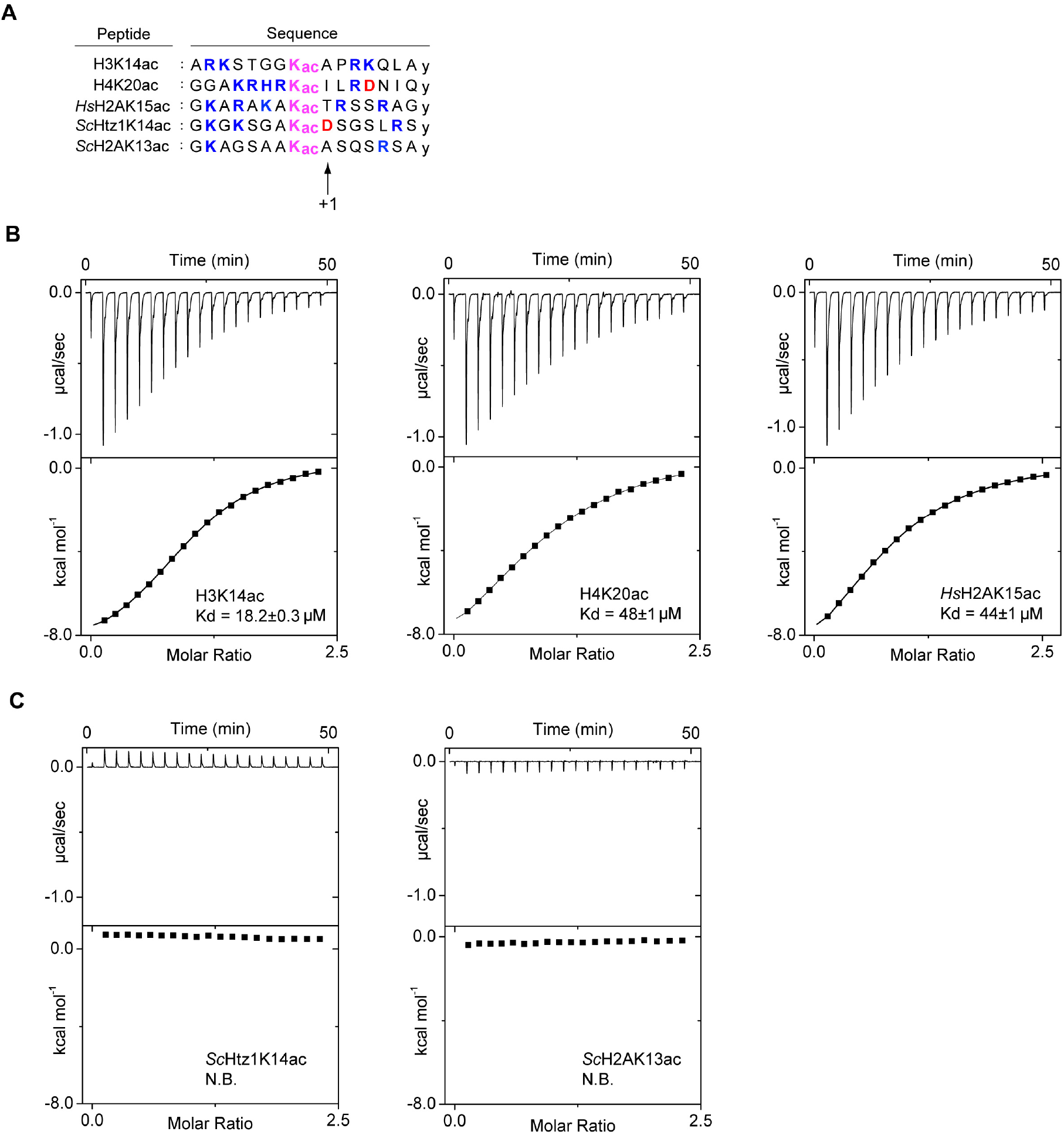
The *Sc*Sth1p Bromodomain Specifically Interacts with Histone Marks Associated with Active Gene Transcription. (A) Primary sequences of the acetylated 15-mer histone peptides used in the ITC binding assay. Acetyl-lysine is highlighted in the center for each peptide. Positively and negatively charged residues are colored in blue and red, respectively. The +1 position in each sequence marks a residue critical for bromodomain specificity. All peptides contain a C-terminal non-histone tyrosine (y) for spectrophotometric concentration determination at 280 nm. (B-C) ITC binding profiles for the *Sc*Sth1p bromodomain interactions with acetylated histone peptides: H3K14ac, H4K20ac, *Hs*H2AK15ac (panel B) and *Sc*Htz1K14ac, *Sc*H2AK13ac (panel C). Raw heat signals generated during each titration are displayed in the top panel as time traces. Below, integrated titration data are displayed and were analyzed by the one-set of sites model (continuous lines). The determined dissociation constants (K_d_s) are displayed for the interacting peptides in (B), whereas no binding (N.B.) was detected in (C).

Inspection of the peptide sequences used in our binding assay reveals that the interacting acetylated histones are enriched in positively charged residues compared to the non-interacting peptides (Figure 2A and Table S1). The available structures of bromodomain complexes with histone peptides indeed indicate that lysine and arginine residues flanking the acetylated lysine participate in electrostatic interactions with the bromodomain (<u>Plotnikov et al., 2014</u>; <u>Zeng et al., 2008</u>). However, when we varied the ionic strength and pH of the buffer, the *K*_d_ values measured by ITC were not significantly altered (up to fourfold) (Figure S2). These results imply that electrostatic forces do not dominate the *Sc*Sth1p bromodomain interactions with the tested acetylated histone peptides.

### The Crystal Structure of the *Sc*Sth1p Bromodomain Reveals a Hydrophobic Acetyl-Lysine Peptide Binding Site

In order to obtain structural insights into histone recognition by the *Sc*Sth1p bromodomain, we determined its crystal structure at 2.7 Å resolution. Crystallographic phases were determined by molecular replacement. The asymmetric unit contains six bromodomains that are highly similar to each other with a root mean square deviation of 0.5 Å. The structure was refined to *R*_work_/*R*_free_ of 23.8%/27.0% and features excellent geometry as evaluated by MolProbity (Table 1) (<u>Chen et al., 2010</u>).

**Table 1.** Data Processing and Refinement Statistics

| Table 1. Data Processing and Refinement Statistics |  |
| --- | --- |
| **PDB ID** | **6MR4** |
| Wavelength [Å] | 0.97918 |
| Space group | C121 |
| Unit cell (*a*, *b*, *c*) [Å] | 141.8, 74.6, 72.5 |
| Unit cell ($\alpha$, $\beta$, $\gamma$) [°] | 90.0, 92.9, 90.0 |
| Resolution [Å] | 72.45-2.71 (2.84-2.71)<sup>a</sup> |
| R<sub>merge</sub> | 0.083 (0.581) |
| <I/σI> | 10.8 (1.7) |
| Completeness [%] | 98.6 (99.5) |
| Redundancy | 3.1 (3.2) |
| CC ½ | 0.99 (0.60) |
| Obs. Reflections | 64,102 (8,612) |
| Unique Reflections | 20,385 (2,722) |
| **Refinement** |  |
| Resolution [Å] | 2.71 |
| No. Reflections | 20,371 |
| R<sub>work</sub>/ R<sub>free</sub> [%] | 23.8/27.0 |
| No. Atoms |  |
| Protein | 5476 |
| Solvent | 27 |
| B-factors [Å<sup>2</sup>] |  |
| Protein | 41.6 |
| Solvent | 58.9 |
| R.m.s. deviations |  |
| Bond lengths [Å] | 0.006 |
| Bond angles [°] | 0.6 |
| All atom clash score | 7.7 |
| Ramachandran<sup>b</sup> [%] |  |
| Favored | 98.74 |
| Allowed | 1.26 |
| Rotamer outliers [%] | 0 |
| C<sub>β</sub> deviation | 0 |

Data were collected on a single crystal.
<sup>a</sup> Values in parentheses are for highest-resolution shell.
<sup>b</sup> As determined by MolProbity.

The *Sc*Sth1p bromodomain folds into a canonical left-handed four-helix bundle (αZ, αA, αB, αC). In the variable ZA loop, the PDY motif contributes to the αAZ helix, while the PMA motif, which deviates from the consensus P*ϕ*D sequence, marks the end of this loop (Figures 1C and 3A) (<u>Dhalluin et al., 1999</u>; <u>Filippakopoulos et al., 2012</u>). Notably, the surface surrounding the acetyl-lysine binding pocket of the bromodomain is relatively hydrophobic, consistent with the lack of large effects on peptide binding affinity at varying salt concentrations and buffer pH (Table S1 and Figure S2). This hydrophobic area distinguishes the *Sc*Sth1p bromodomain from the polar site of its human homolog *Hs*SMARCA2 (Figure S4).

### The *Sc*Sth1p Bromodomain Contains a β-Hairpin Characteristic of Family VIII Members

The β-hairpin located between the αZ and αZ’ helices classifies the *Sc*Sth1p bromodomain as a family VIII member (Figures 1C and 3B). A distance matrix alignment (DALI) search (<u>Holm and Rosenstrom, 2010</u>) against known structures in the Protein Data Bank (PDB) indeed confirms that the *Sc*Sth1p bromodomain is structurally most similar to other family VIII members, including bromodomains of *Hs*SMARCA2 and *Hs*PB1 (Figures 1C and 3B) (<u>Filippakopoulos et al., 2012</u>; <u>Singh et al., 2007</u>)). The β-hairpin in the *Sc*Sth1p bromodomain is stabilized by a salt bridge between His1272 and Asp1342 in the αC helix (Figure 3C). Disruption of this salt bridge by the His1272Ala mutation did not affect peptide specificity or binding affinity, suggesting that it does not significantly contribute to recognizing acetylated histones (Figures 3D and S3A).

**Figure 3.**
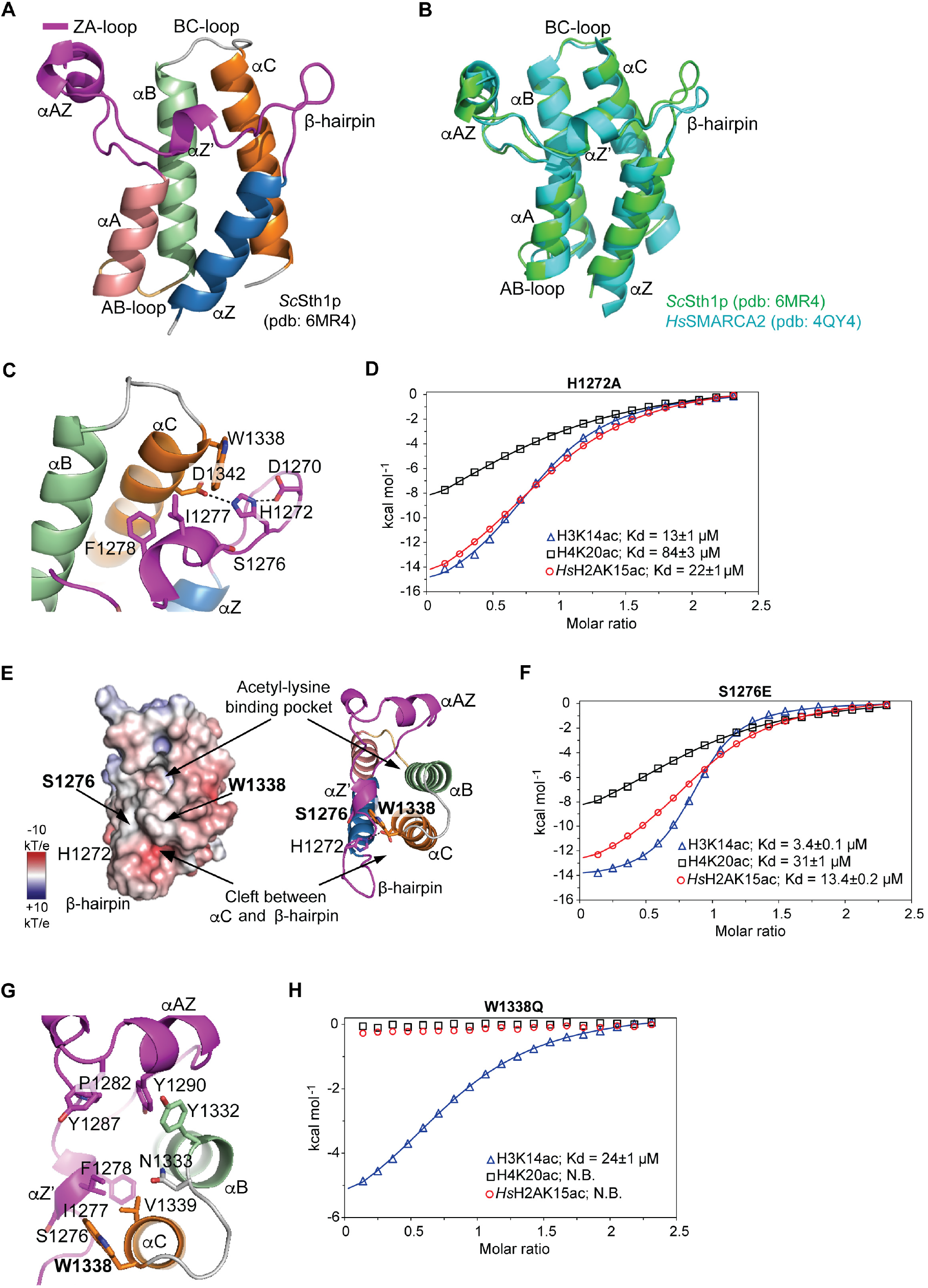
Structure and Mutational Analysis of the *Sc*Sth1p Bromodomain. (A) Overall structure of the *Sc*Sth1p bromodomain shown with labeled secondary structure elements: αZ (blue), ZA-loop (magenta), αB (green), αC (orange) and αA (pink). The ZA-loop contains the SIF motif as part of the αZ’ helix and the β-hairpin. (B) Structural alignment of the *Sc*Sth1p and *Hs*SMARCA2 (pdb: 4QY4) bromodomains. (C) A detailed view of the interactions between the β-hairpin and αC helix. His1272 in the β-hairpin forms two hydrogen bonds: one with Asp1270 in the β-hairpin, the other with Asp1342 in the αC helix. (D) ITC binding profiles for interactions between the *Sc*Sth1p His1272Ala mutant bromodomain and H3K14ac, H4K20ac or *Hs*H2AK15ac peptides. Binding data were analyzed by the one-set of sites model (continuous lines) and the obtained K_d_ values are displayed. (E) Electrostatic surface potential of the *Sc*Sth1p bromodomain (left panel). Positive and negative surface potential is colored in blue and red, respectively. The electrostatic surface potential scale is shown, where k_B_ is the Boltzmann constant and T is the absolute temperature. A cartoon representation of the *Sc*Sth1p bromodomain structure is displayed in the same orientation with the mutated residues highlighted (right panel). The hydrophobic acetyl-lysine binding pocket and the slightly negatively charged cleft between the β-hairpin and αC helix are indicated. (F) ITC binding profiles for the interactions between the *Sc*Sth1p Ser1276Glu mutant bromodomain and H3K14ac, H4K20ac, and *Hs*H2AK15ac peptides. Binding data were analyzed by the one-set of sites model (continuous lines) and the obtained K_d_ values are displayed. (G) A close-up view of the acetyl-lysine binding site in the *Sc*Sth1p bromodomain. (H) ITC binding profiles for the interactions between the *Sc*Sth1p Trp1338Gln mutant bromodomain and H3K14ac, H4K20ac, and *Hs*H2AK15ac peptides. The H3K14ac peptide binding data were analyzed by the one-set of sites model (continuous lines), whereas the remaining peptides do not bind (N.B.).

### Two Single Point Mutations in the *Sc*Sth1p Bromodomain Dramatically Increase Substrate Affinity and Specificity

Another common feature of the family VIII bromodomains is the E/D*ϕ*F sequence, which substitutes for the “WPF shelf motif” found in many bromodomains (Figures 1C and 3A). However, the corresponding “SIF” motif in *Sc*Sth1p deviates from this consensus sequence. Like the WPF motif, the SIF sequence folds into a short 3_10_-helix termed αZ’. To examine the contribution of this non-conserved serine to acetylated histone peptide binding (Figure 3E), we mutated it to the more common glutamate found in several family VIII bromodomains (Figure 1C). Strikingly, the Ser1276Glu mutation increases peptide binding affinity by up to sixfold with respect to the wild-type bromodomain, resulting in a tenfold tighter binding to H3K14ac than to H4K20ac (K_d_ values of 3 and 31 µM, respectively; Figures 3F and S3B).

In addition to classifying bromodomains based on their entire structures (<u>Filippakopoulos et al., 2012)</u>, they can also be grouped according to the conservation of signature residues in their variable acetyl-lysine binding sites (<u>Vidler et al., 2012</u>). According to the latter classification, the *Sc*Sth1p bromodomain shares conserved elements with classes 5 and 6. These include residues Ser1337, Val1339, Asp1342, and Tyr1287 that line the acetyl-lysine binding pocket (Figures 3G and 1C). Notably, Trp1338 at the end of the BC loop is not present in any other bromodomain and the equivalent position is most often occupied by Ala, Glu, or Gln (<u>Filippakopoulos et al., 2012</u>; <u>Zhang et al., 2010</u>). When we mutated Trp1338 to the more common glutamine found in family VIII members (Figure 1C), we observed a dramatic effect on substrate specificity. While the Trp1338Glu mutant bromodomain binding to H3K14ac was comparable to the wild-type protein, its interactions with the H4K20ac and *Hs*H2AK15ac peptides were completely abolished (Figures 3H and S3C). Hence, our results show that two single point mutations of the two non-conserved residues, Ser1272 and Trp1338, significantly increase *Sc*Sth1p bromodomain binding affinity and specificity towards acetylated histone peptides.

## Discussion

One of the major challenges in chromatin biology is to understand how distinct patterns of post-translational modifications are recognized and “interpreted” or “read” by chromatin interacting proteins and complexes. Bromodomains are ubiquitous protein modules that bind site-specific acetylated lysines in histones, a key modification implicated in transcriptional activity. Most bromodomains recognize multiple acetylated substrates with a wide range of selectivity and modest binding affinity (<u>Filippakopoulos and Knapp, 2012</u>). While it has been accepted that such binding behavior can be understood in light of cooperative engagement with several histone modifications (<u>Ruthenburg et al., 2007</u>), the question arises whether there are intrinsic, evolutionary limitations to the bromodomain fold that preclude it from recognizing substrates with high, i.e. nanomolar affinities. To address this issue, we focused on the *Sc*Sth1p bromodomain, which binds several acetylated histone peptides with apparent high affinity in a dot-blot assay (<u>Zhang et al., 2010</u>). We discovered that it was surprisingly straightforward to substantially increase both substrate affinity and specificity by generating two point mutations in the bromodomain, changing only one residue at a time.

The first of these two key residues is Ser1276 of the SIF motif within the variable ZA loop, which deviates from the acidic residues in the consensus E/D*ϕ*F sequence of family VIII bromodomains and from the tryptophan in the WPF motif present in several other bromodomain families (Figure 1C) (Filippakopoulos et al., 2012). For example, Ser1276 corresponds to Glu206 in the second bromodomain of the human polybromodomain protein PB1, the only family VIII member whose structure has been determined in complex with a histone peptide to date (<u>Charlop-Powers et al., 2010</u>). In this NMR structure, Glu206 plays a key role in recognizing the Kac +1 residue alanine (Ala15) of the H3K14ac peptide, discriminating histone peptides with a sequence variation at this position (<u>Charlop-Powers et al., 2010</u>). Consistent with this observation, the Ser1276Glu mutation largely increased selectivity for H3K14ac over H4K20ac or *Hs*H2AK15ac peptides. We envision that the longer side chain of Glu1276 in the mutant bromodomain is able to accommodate the Ala side chain at the +1 position in H3K14ac, but the bulkier side chains of Ile and Thr in the H4K20ac or *Hs*H2AK15ac peptides fit to a lesser degree. We conclude that the exposed ZA loop residue at the position equivalent to Ser1276 is a major determinant of substrate affinity in the family VIII bromodomains.

The second residue that we identified as a key determinant for the *Sc*Sth1p bromodomain-peptide interactions corresponds to Trp1338 at the end of the BC loop, proximal to Ser1276 (Figure 3E). Alignment of the 61 human and 14 yeast bromodomain sequences illustrates that a tryptophan residue is unique at this position, which is occupied by a glutamine in the majority of family VIII members (<u>Filippakopoulos et al.,2012</u>). Strikingly, our Trp1338Gln mutation had an even larger impact on peptide recognition than the Ser1276Glu mutation; it completely abolished the *Sc*Sth1p bromodomain binding to the H4K20ac and *Hs*H2AK15ac peptides, while the affinity for H3K14ac essentially remained unchanged. These results suggest that the large aromatic surface area of the exposed Trp1338 may contribute extensive contacts with the interacting peptides.

Our proposed role for Trp1338 is corroborated by its prominent location at the intersection of two roughly perpendicular grooves that are poised to accommodate bound peptides. The first groove is hydrophobic and extends above the canonical N-acetyl-lysine binding pocket between the ZA and BC loops, which corresponds to the preferred peptide binding groove in experimentally determined bromodomain-peptide complexes (Figure 3E) (<u>Fujisawa and Filippakopoulos, 2017</u>; <u>Marchand and Caflisch, 2015</u>; <u>Zeng et al., 2008</u>). The second groove is slightly negatively charged and is formed between the β-hairpin and a surface patch formed by Trp1338 and Asp1267. A binding mode of peptides wrapping around Trp1339 would be consistent with the large effect that the Trp1339Gln mutation has on peptide recognition. Precedence for such a binding mode is found in the structure of the *Hs*Brd4 bromodomain complex with a diacetylated histone peptide (PDB code 3UVW), in which the peptide bends around Asp145 at the Trp1339-equivalent position at a 90° angle (<u>Filippakopoulos et al., 2012</u>). In contrast to Trp1338, the opposite edge of the second groove appears to be less important for peptide recognition, as the His1272Ala mutation, which presumably destabilizes the β-hairpin, has a negligible effect on peptide binding. Overall, the predominantly apolar character of the two binding grooves agrees well with our finding that the *Sc*Sth1p bromodomain-peptide interactions are dominated by hydrophobic contacts.

Our results on substrate specificity of the *Sc*Sth1p bromodomain have important implications for the recruitment of its resident RSC chromatin remodeling complex. In *S. cerevisiae*, the H3K14ac and Htz1K14ac marks are enriched at TSSs, but it is unknown whether both are required for chromatin remodeling initiation (<u>Pokholok et al., 2005</u>; <u>Ramachandran et al., 2015</u>). As the *Sc*Sth1p bromodomain recognizes H3K14ac, but not the *Sc*Htz1K14ac mark (Figure 2C), our data suggest that *Sc*Sth1p recruits the RSC complex to TSSs via the interaction with the H3K14ac mark. Indeed, Htz1-depleted cells do not alter nucleosome positioning at TSSs in yeast (<u>Hartley and Madhani, 2009</u>), further suggesting that the Htz1 histone H2A variant may not be required for RSC recruitment to chromatin. Given that *Sc*Rsc4p represents another H3K14ac reader module in the RSC complex (<u>Kasten et al., 2004</u>; <u>VanDemark et al., 2007</u>), these two essential RSC components may cooperatively bind to two histone tails, thereby increasing the binding affinity and specificity of the RSC complex at active TSSs (Figure 4). In addition to the H3K14ac binding, the *Sc*Sth1p bromodomain recognizes the H4K20ac modification, yet its biological function in yeast remains elusive.

**Figure 4.**
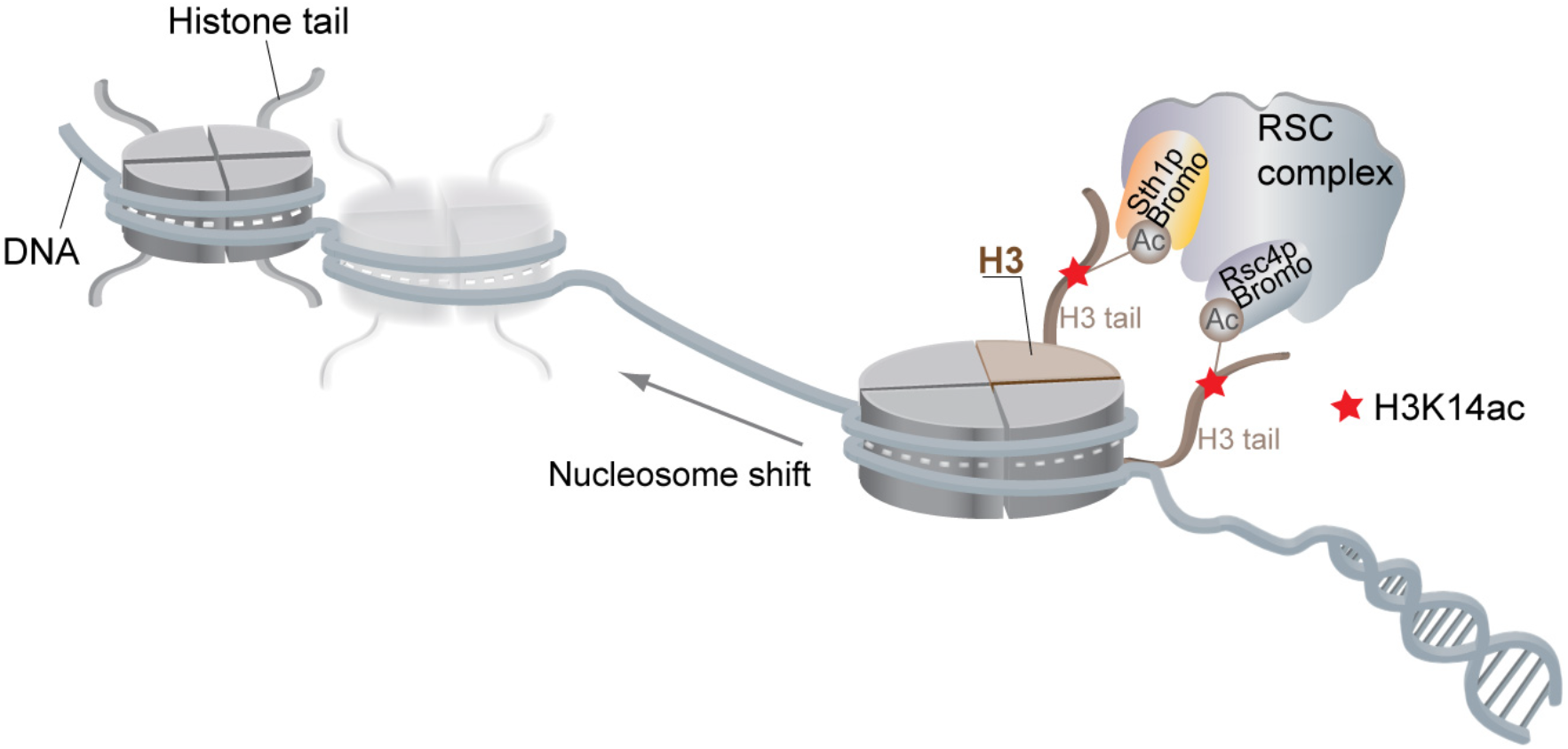
The *Sc*Sth1p Bromodomain Contributes to the RSC Complex Association with H3K14ac-Modified Chromatin. A proposed schematic representation of the RSC complex binding to transcriptionally active chromatin. The bromodomains in *Sc*Sth1p and *Sc*Rsc4p—key members of the RSC complex—are thought to interact with acetylated histones to direct RSC to promoter transcription start sites (TSSs) marked with H3K14 acetylation. The remaining bromodomain-containing components of the RSC complex, *Sc*Rsc1 and *Sc*Rsc2, are omitted for clarity. Once attached to promoters, the RSC complex creates a nucleosome-free region flanked by strongly positioned nucleosomes.

In conclusion, our structure-based mutational analysis shows that substrate binding affinity and specificity are finely balanced by individual residues in the *Sc*Sth1p bromodomain. Instead of recognizing a single chromatin mark with high affinity, our results are consistent with the notion that multiple histone modifications—each interacting with intermediate affinity—are cooperatively “read” with high affinity and specificity. In this scenario, the bromodomains of the RSC components *Sc*Sth1p and *Sc*Rsc4p may cooperatively facilitate recruitment of the RSC complex towards a subset of TSSs marked by the H3K14 acetylation. Our *Sc*Sth1p bromodomain point mutants may prove as valuable tools to probe the mechanism of *Sc*Sth1p-dependent recruitment of the RSC complex to chromatin *in vivo*.

## Materials and Methods

### Protein Expression and Purification

The DNA fragment corresponding to the *S. cerevisiae* Sth1p bromodomain (residues 1250-1359) was generated by PCR from genomic DNA and cloned into a modified pGEX-4T1 vector (Addgene) using Ligation Independent Cloning (LIC), as previously described (<u>Seo et al., 2013</u>). The resulting fusion protein contained an N-terminal GST tag followed by a cleavage site for the tobacco etch virus (TEV) protease. The *Sc*Sth1p bromodomain was expressed in *E. coli* BL21-CodonPlus(DE3)-RIL cells (Stratagene) following overnight induction at an OD_600nm_ of 0.8 with 0.5 mM isopropyl-β-D-thio-galactoside (IPTG) at 18°C in LB media containing chloramphenicol (34 mg/L) and ampicillin (100 mg/L). Cells were lysed using a cell disruptor (Avestin) in a lysis buffer containing 20 mM HEPES (pH 7.5), 300 mM NaCl, 0.5 mM EDTA, 5% (v/v) glycerol, 3 mM DTT, 1 mM benzamidine, supplemented with phenylmethanesulfonyl fluoride (PMSF), 1 µg/mL bovine lung aprotinin and leupeptin (Sigma). After centrifugation at 30,000 × *g,* the supernatant was loaded onto a GSTrap column (GE Healthcare) and washed extensively with lysis buffer. The *Sc*Sth1p bromodomain was eluted with 20 mM HEPES pH 7.5, 200 mM NaCl, 5% (v/v) glycerol, and 3 mM DTT supplemented with 15 mM reduced glutathione (Sigma). Following an on-column GST-tag cleavage by TEV protease at 4°C overnight, the bromodomain was further purified over a HiLoad 16/60 Superdex 75 gel filtration column (GE Healthcare) equilibrated with 20 mM HEPES pH 7.5, 200 mM NaCl, and 3 mM DTT. Peak elution fractions were concentrated to 5-10 mg/mL, flash-frozen in liquid nitrogen, and stored at −80°C.

### Peptide Synthesis

Synthetic histone peptides for ITC binding experiments were synthesized and HPLC-purified to 95-98% purity at the Proteomics Resource Center (The Rockefeller University). The correct sequence for each peptide was confirmed by mass spectrometry. All peptides are 15 amino acids long and contain a C-terminal non-histone tyrosine for concentration measurements at OD_280nm_.

### *Sc*Sth1p Bromodomain Mutagenesis

Constructs for the *Sc*Sth1p bromodomain mutants (Ser1276Glu, His1272Ala, Trp1338Gln, or Tyr1290Ala/Asn1333Ala) were generated using QuikChange site-directed mutagenesis kit (Stratagene). Mutant bromodomain expression and purification were carried out as for the wild-type protein. Gel filtration-purified protein was concentrated to 5-10 mg/mL, flash-frozen in liquid nitrogen and stored at −80°C.

### ITC Binding Assays

Samples for the ITC experiments were dialyzed extensively against 20 mM HEPES pH 7.5, 150 mM NaCl, 0.5 mM TCEP using Float-A-Lyzer dialysis devices with a molecular weight cutoff of 0.5-1 kDa (Spectrum Laboratories). For a subset of ITC experiments, composition of the binding buffer (pH 7.5) was adjusted to 75 or 300 mM NaCl, or the buffer pH was set to 6.0 or 8.5 at 150 mM NaCl (see Table S1 for details). After dialysis, purified bromodomain and histone peptides were filtered (0.22 µm) and centrifuged, followed by determining their concentration by UV absorbance at 280 nm. All titration experiments were performed using an automated MicroCal Auto-iTC200 instrument (GE Healthcare) at 15°C. For each titration, a concentrated solution of the peptide (at 1 mM) was titrated into the ITC reaction cell containing the wild-type or mutant *Sc*Sth1p bromodomain (at 90 µM). The heat of sample dilution was separately measured for each experiment or determined from the final plateau of each titration, and subtracted from the binding data. The corrected ITC curves were analyzed using a nonlinear least-squares minimization method in ORIGIN 7.0 to determine the binding stoichiometry (n), dissociation constant (K_d_) and enthalpy (ΔH). These parameters were subsequently used to determine free Gibbs energy (ΔG) and the entropic component (TΔS) using ΔG = −RT ln(1/K_d_) and TΔS = ΔH – ΔG equations, where R and T are gas constant (1.99 cal/(mol*K)) and absolute temperature (288 K), respectively. The fitting parameters were allowed to float freely in the fitting procedure. All titrations were repeated at least twice. Thermodynamic parameters for representative ITC titrations are provided in Table S1.

### Crystallization and Structure Determination of the *Sc*Sth1p Bromodomain

Crystals of the *Sc*Sth1p bromodomain were grown at 20°C in sitting drops containing 1 µL of the protein at 5 mg mL^-1^ and 1 µL of a reservoir solution consisting of 0.2 M magnesium chloride hexahydrate, 0.1 M BIS-TRIS pH 6.5, 25% (w/v) polyethylene glycol 3,350, and 3% (w/v) trimethylamine N-oxide dihydrate. For cryoprotection, crystals were stabilized in the mother liquor supplemented with 20% (v/v) glycerol and flash cooled in liquid nitrogen. X-ray diffraction data were collected at 100 K at the NE-CAT beamline 24ID-E at the Advanced Photon Source, Argonne National Laboratory, and processed using XDS (<u>Kabsch, 2010</u>). The crystals belong to the space group C121, with unit cell dimensions of a=141.8 Å, b=74.6 Å, c=72.5 Å, α=γ=90°, and β=92.9° and diffracted to 2.7 Å resolution (Table 1). Crystallographic phases were obtained by molecular replacement using the fourth bromodomain of human poly-bromodomain containing protein 1 (PB1; pdb: 3TLP) as a search model using PHASER (<u>McCoy et al.,2007</u>) from the CCP4 program suite. After molecular replacement, iterative cycles of manual rebuilding and restrained refinement were carried out in COOT (<u>Emsley and Cowtan, 2004</u>) and PHENIX (<u>Adams et al., 2010</u>), respectively. The stereochemical quality of the structural model was assessed with MolProbity (<u>Chen et al., 2010</u>). There were no residues in the disallowed region of the Ramachandran plot (Table 1). Figures were generated with PyMOL (DeLano Scientific) (<u>Schrodinger, 2015</u>).

## Acknowledgements

First and foremost, we would like to acknowledge our mentor G. Blobel for supporting our research. We would like to thank C. Adura for helping us with the Auto-iTC200 instrument (High-Throughput and Spectroscopy Resource Center, Rockefeller University), H. Zebroski III for peptide synthesis (Proteomics Resource Center, Rockefeller University), and R. Rajashankar (NE-CAT) for assisting with x-ray data collection. We would like to thank E. Coutavas for critical reading of the manuscript. This work was funded by the Rockefeller University and the Howard Hughes Medical Institute.

## Author Contributions

B.J.B., H.H, H-S.S, and E.W.D designed the experiments; B.J.B., H.S-S., and A.K performed the experiments; B.J.B., H.H., H-S.S, and E.W.D analyzed the data, and B.J.B., H.H., A.K., and E.W.D. wrote the manuscript.

## Accession Code

The *Sc*Sth1p bromodomain coordinates and structure factors have been deposited at the Protein Data Bank (PDB) with the accession number 6MR4.

## Supplemental Information

**Figure S1.**
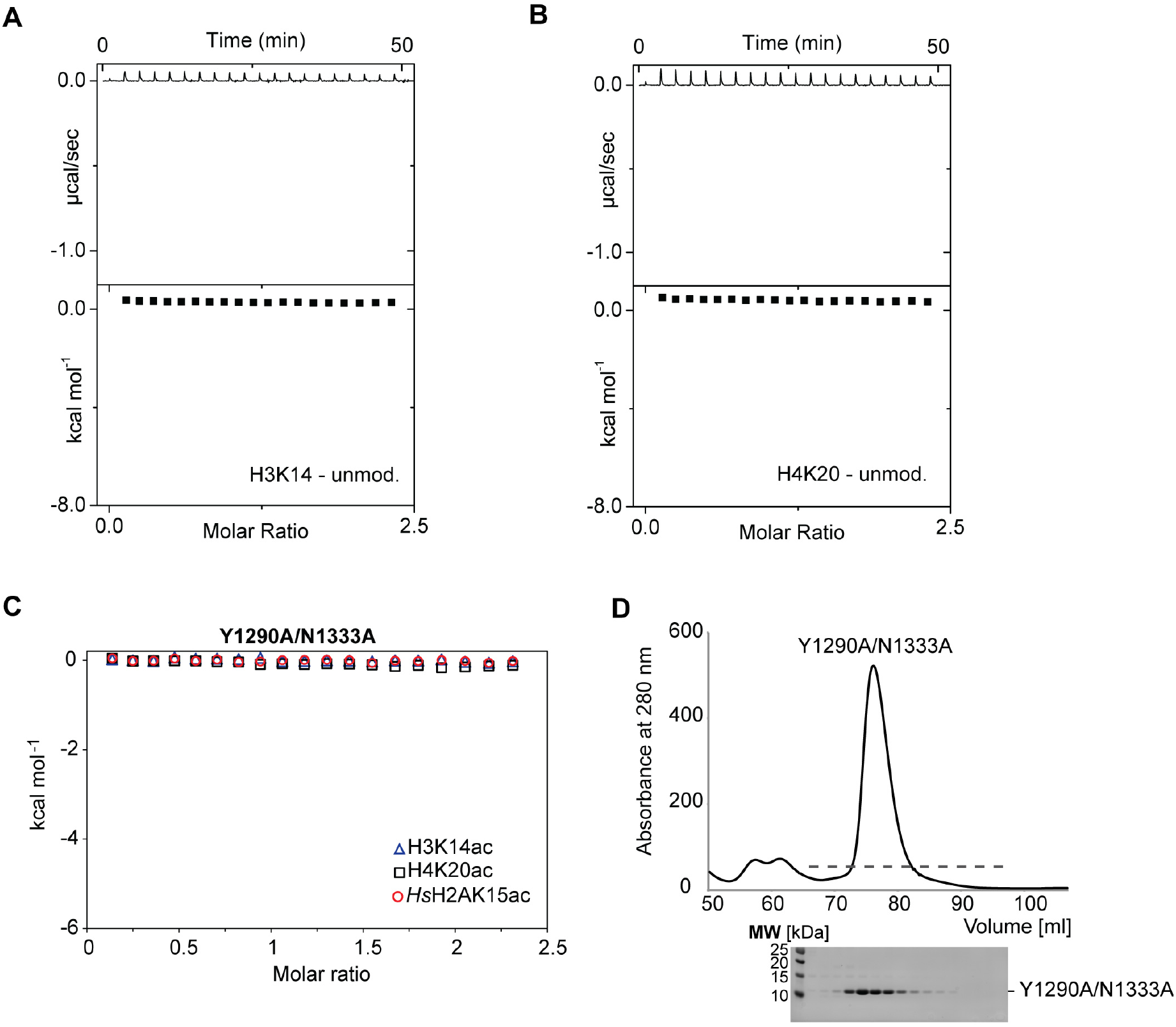
Related to Figure 2. The *Sc*Sth1p Bromodomain Contains a Canonical Binding Pocket for Acetyl-Lysine Recognition. (**A-B**) ITC binding profiles for the *Sc*Sth1p bromodomain interactions with unmodified (non-acetylated) (**A**) H3K14 and (**B**) H4K20 peptides are shown in the top panel as time traces. No appreciable heat of binding (N.B.) was observed for either peptide. (**C**) ITC binding profiles for the *Sc*Sth1p Tyr1290Ala/Asn1333Ala mutant bromodomain interactions with H3K14ac, H4K20ac, and *Hs*H2AK15ac peptides. No appreciable heat of binding (N.B.) was observed. (**D**) A size-exclusion chromatography (SEC) elution profile for the *Sc*Sth1p bromodomain mutant bromodomain (Tyr1290Ala/Asn1333Ala). Below, peak elution fractions were resolved by SDS-PAGE. The mutant *Sc*Sth1p bromodomain elutes as a single monomeric peak.

**Figure S2.**
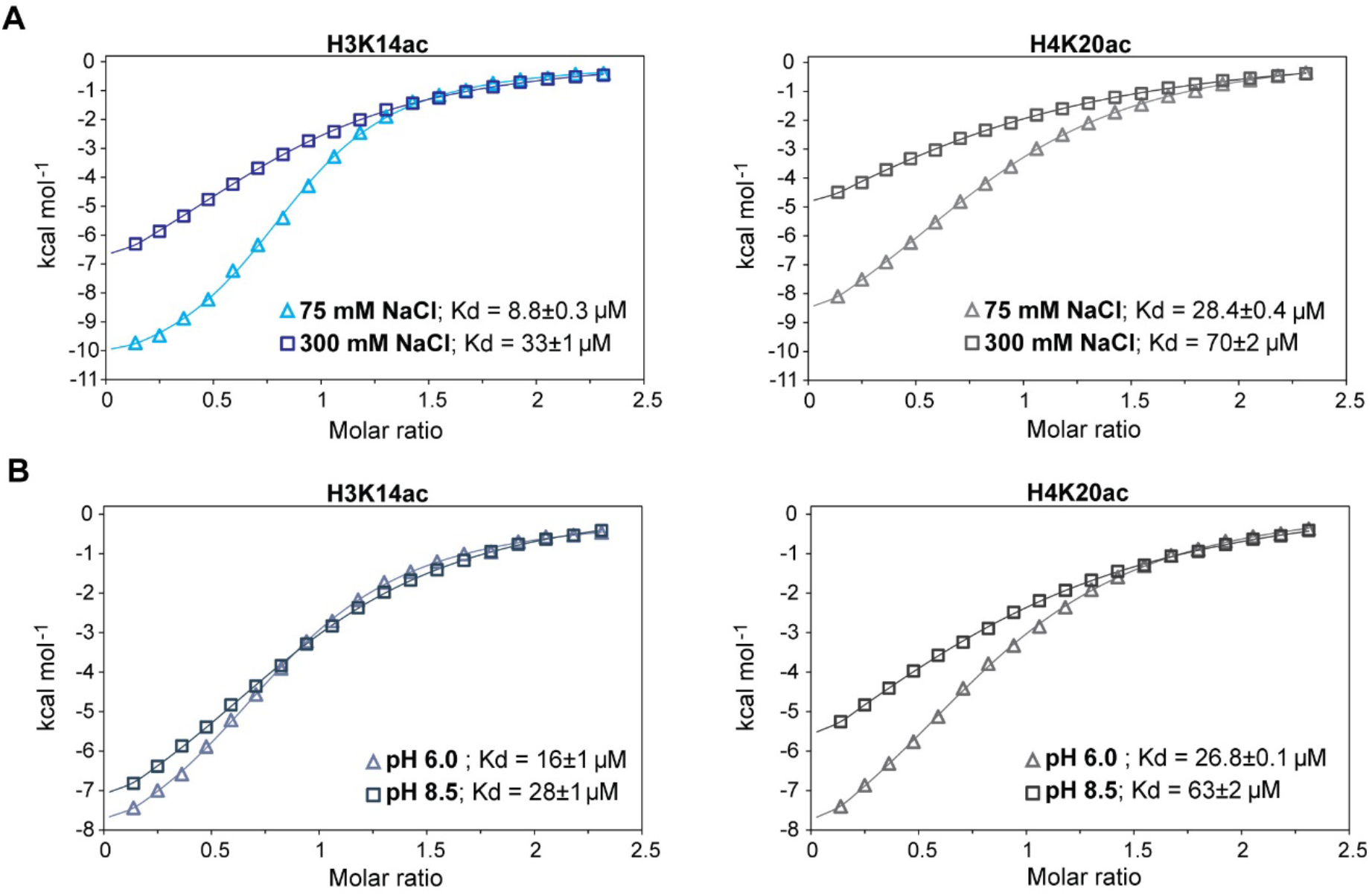
Related to Figure 2. Effect of Buffer Ionic Strength and pH on *Sc*Sth1p Bromodomain Interactions with Acetylated Histone Peptides. (**A-B**) ITC binding profiles for the *Sc*Sth1p bromodomain interactions with H3K14ac and H4K20ac peptides at various binding conditions. ITC titrations were performed in 20 mM HEPES pH 7.5, 0.5 mM TCEP supplemented with 75 mM or 300 mM NaCl (panel A) or in 20 mM HEPES pH 6.0 or 8.0, 0.5 mM TCEP, and 150 mM NaCl (panel B). Binding data were analyzed by the one-set of sites model (continuous lines) and the obtained K_d_ values are displayed.

**Figure S3.**
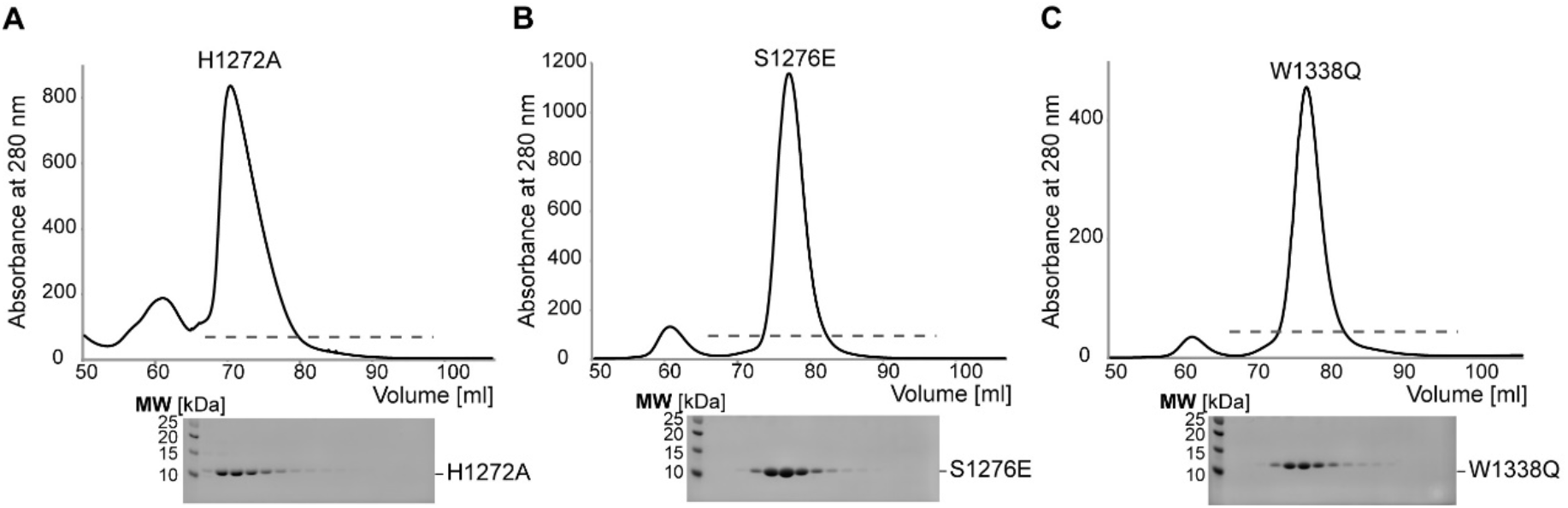
Related to Figure 3. SEC Elution Profiles for the *Sc*Sth1p Mutant Bromodomains. (**A-C**) Size-exclusion chromatography (SEC) elution profiles for the *Sc*Sth1p bromodomain mutants. (**A**) The *Sc*Sth1p His1272Ala mutation likely affects the stability of the β-hairpin, resulting in a shifted and asymmetric elution profile. (**B-C**) Ser1276Glu and Trp1338Gln mutant bromodomains elute like the wild-type protein. Below, peak elution fractions were resolved by SDS-PAGE.

**Figure S4.**
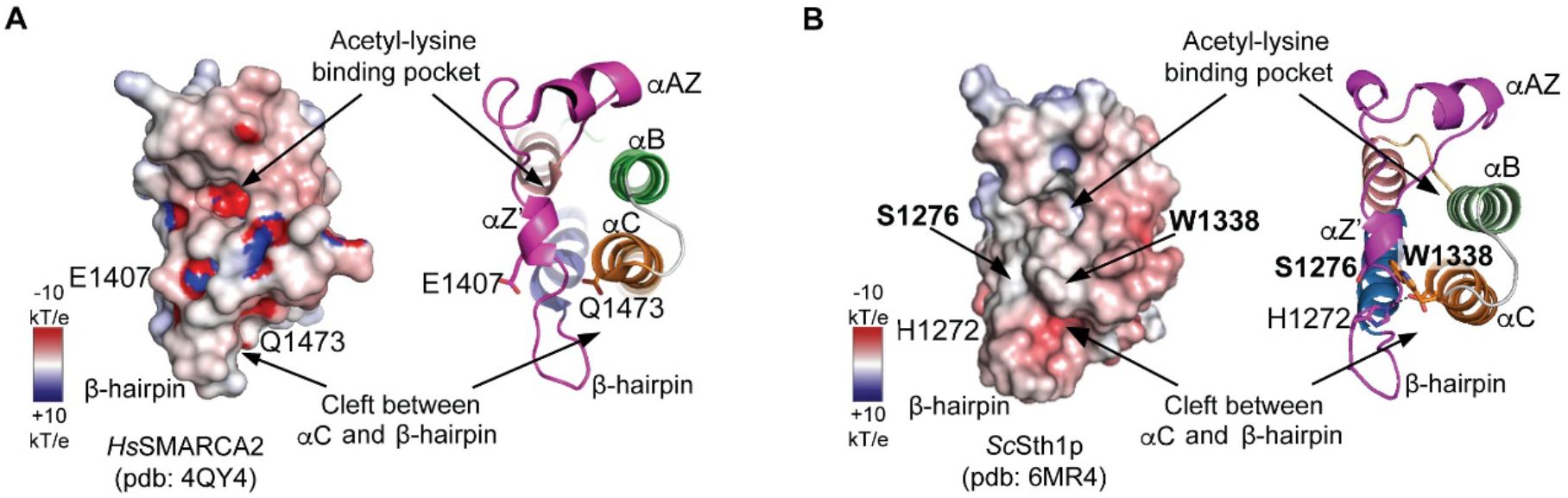
Related to Figure 3. Surface Charge Distribution of the *Sc*Sth1p and *Hs*SMARCA2 Bromodomains. (**A**) Surface charge distribution of the human SMARCA2 bromodomain (left panel). Positively charged regions of the structure are colored in blue, whereas negative and neutral regions are labeled in red and white, respectively. The electrostatic potential scale is shown, where k_B_ is the Boltzmann constant and T stands for absolute temperature. The structure is displayed as a cartoon in the same view (right panel). The negatively-charged acetyl-lysine binding pocket and the hydrophobic cleft formed by the β-hairpin and αC helix are highlighted. These structural features distinguish the *Hs*SMARCA2 and *Sc*Sth1p bromodomains (**B**). The highlighted Glu1407 and Gln1473 residues in *Hs*SMARCA2 (**A**) correspond Ser1276 and Trp1338 in the *Sc*Sth1p bromodomain (**B**), which we mutated in this study.

**Table S1:**
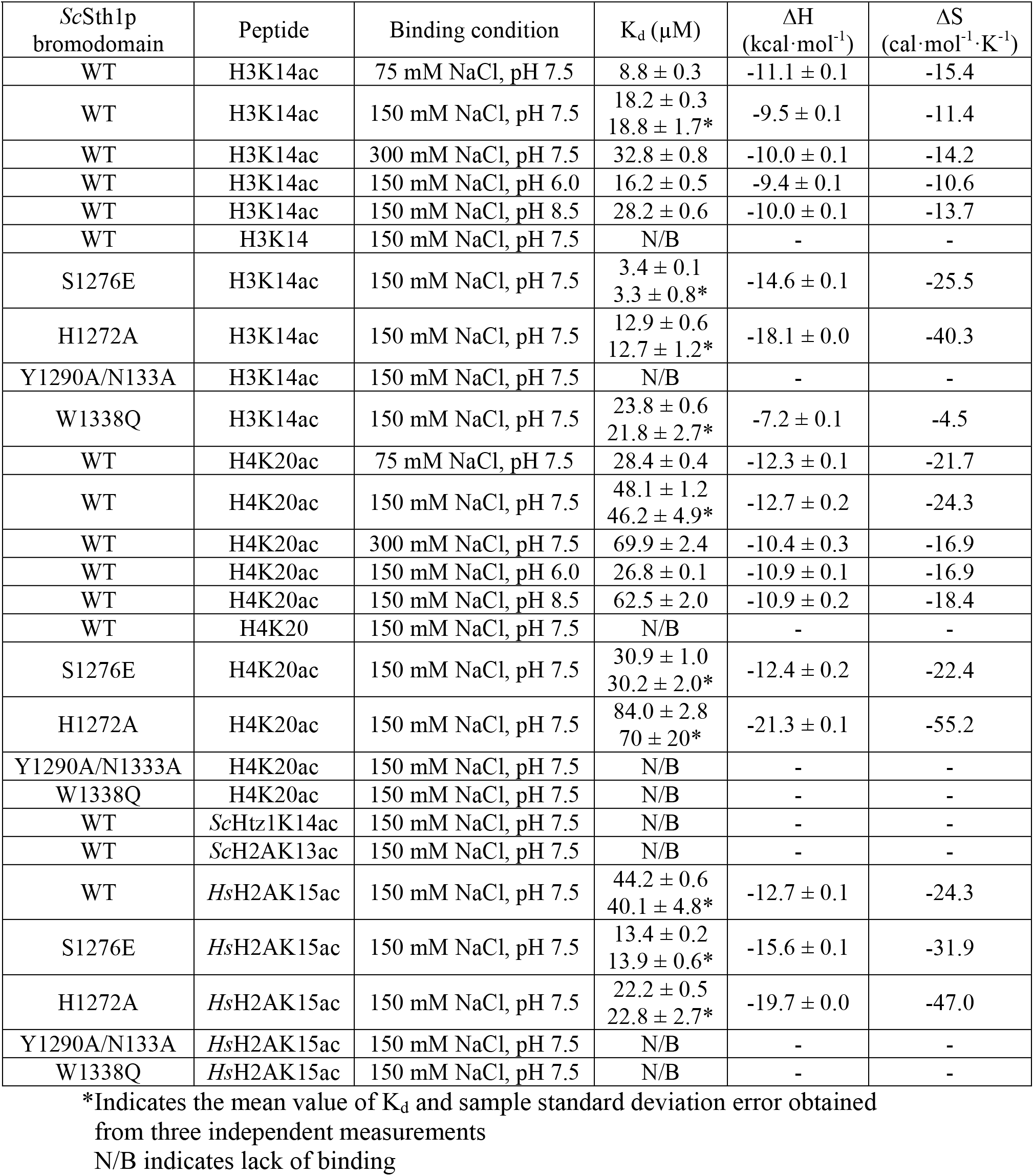
Thermodynamic Parameters for the *Sc*Sth1p Bromodomain Binding to Acetylated Histone Peptides

